# Tubulin Monoglutamylation is Sufficient to Rescue the Ciliary Motility Defects in a *Chlamydomonas* Polyglutamylation Deficient Mutant

**DOI:** 10.64898/2026.04.09.717589

**Authors:** Rinka Sasaki, Toshiyuki Oda, Tomohiro Kubo

**Affiliations:** Department of Anatomy and Structural Biology, Graduate School of Medicine, University of Yamanashi, 1110 Shimokato, Chuo, Yamanashi, 409-3898, Japan

**Author notes:** Corresponding authors, Email addresses. **Author Contributions:** T.K. conceived and designed experiments. R.S. and T.K. performed the experiments and analyzed the data. R.S. and T.K. drafted the article. T.K. and T.O. acquired funding and supervised the project.

**Keywords:** Cilia, tubulin, glutamylation, tubulin tyrosine ligase-like (TTLL) protein, cytosolic carboxypeptidase (CCP), *Chlamydomonas reinhardtii*

## Abstract

The axonemes of eukaryotic cilia and flagella display high tubulin glutamylation heterogeneity, yet the functional significance of this variation remains elusive. We previously showed that long-chain polyglutamylation is crucial for ciliary motility in *Chlamydomonas*. However, the respective contributions of long-chain polyglutamylation versus short-chain species to motility remain unclear, as existing mutants did not allow for a clear functional dissection of these two modification states. Here, we generated mutants deficient in deglutamylases, cytosolic carboxypeptidases (CCPs) 1, 2, and 5. Importantly, CCP5 is known to remove the branch-point glutamate residue, the final step in deglutamylation. While axonemal polyglutamylation levels remained largely unaffected in these mutants, abundance of short-chain glutamylation was significantly increased in both the axonemal and cytoplasmic microtubules of *ccp5-1*, consistent with CCP5’s role as a branch-point deglutamylase. Although each single mutant exhibited slightly reduced swimming velocity, the loss of *CCP5* in the *tpg1* background lacking long polyglutamate side chains resulted in a significant restoration of motility. These findings indicate that the abundance of short-chain species, regulated by CCP5, plays a distinct role in modulating ciliary motility, particularly in the absence of long polyglutamate side chains. This suggests that even minimal glutamylation can functionally support dynein-driven microtubule sliding.

## INTRODUCTION

α- and β-tubulin undergo diverse post-translational modifications (PTMs), including acetylation, tyrosination, glutamylation, glycylation, methylation, and phosphorylation (Westermann and Weber, 2003; Janke and Bulinski, 2011). Tubulin PTMs modulate the microtubule dynamics and stability, as well as their interactions with various microtubule-associated proteins. Given that microtubule dysfunction caused by PTM defects is linked to human diseases (Magiera et al., 2018; McKenna et al., 2023), understanding the biochemical impact of these PTMs is crucial not only for elucidating the fundamental properties of microtubule cytoskeleton, but also for understanding its critical role in human health.

Cilia (often used interchangeably with flagella) are whip-like organelles essential for cellular motility, fluid transport, and signal transduction. The axoneme, the core structure of cilia, consists of highly stable microtubules that accumulate a high density of diverse tubulin PTMs. Acetylation of α-tubulin was first reported in *Chlamydomonas* cilia (L’Hernault and Rosenbaum, 1985) with its modification site subsequently identified as lysine 40 (LeDizet and Piperno, 1987). Detyrosination, the removal of the C-terminal tyrosine residue from α-tubulin, was later reported in cilia by Stephens (1992). Subsequent studies revealed a striking spatial segregation of these modifications within the axoneme; for instance, Multigner et al. (1996) biochemically demonstrated that detyrosinated tubulin is enriched in the B-tubules of outer doublets, a localization later confirmed morphologically by Johnson (1998). Furthermore, large polymeric modifications on the tubulin C-terminal regions have been identified. Eddé et al. (1990) discovered polyglutamylation, which later was found to be highly abundant in cilia across diverse species. Similarly, Redeker et al. (1994) identified polyglycylation as another major polymeric modification characteristic of axonemal microtubules.

Among these PTMs, polyglutamylation is particularly crucial for its regulation in the ciliary motility (Kubo et al., 2010; Suryavanshi et al., 2010; Ikegami et al., 2010).

The enzymes involved in tubulin polyglutamylation have been extensively characterized. Members of the tubulin tyrosine ligase-like (TTLL) protein family (Janke et al., 2005) catalyze the addition of polyglutamate side chains to the tubulin tail region. Specifically, TTLL1, 4, 5, and 7 function as initiating enzyme, while TTLL6, 7, 9, 11, and 13 are primarily responsible for chain elongation (Ikegami et al., 2006; van Dijk et al., 2007; Wloga et al., 2008; Mukai et al., 2009; Kubo et al., 2010). On the other hand, deglutamylation is mediated by the cytosolic carboxypeptidase (CCP) family. Within this group, CCP1, 2, 3, 4, 6 catalyze the shortening of the glutamate chains (Rogowski et al., 2010; Tort et al., 2014). In contrast, CCP5 uniquely targets the branch-point glutamate to fully remove the modification (Rogowski et al., 2010). Through the action of these enzymes, polyglutamate side chains of diverse lengths are synthesized (Eddé et al., 1990; Alexander et al., 1991; Redeker et al., 2005), providing structural and functional complexity to the microtubule cytoskeleton.

The *tpg1* mutant of *Chlamydomonas reinhardtii* deficient in TTLL9 has provided a valuable model for studying tubulin glutamylation (Kubo et al., 2010; 2012; 2017). The loss of long polyglutamate side chains on axonemal α-tubulin leads to impaired ciliary motility in *tpg1* (Kubo et al., 2010). We previously demonstrated that these side chains regulate the interaction between the adjacent outer-doublet microtubules by modulating the function of the nexin-dynein regulatory complex (N-DRC; Kubo et al., 2012; 2017). However, while the role of long polyglutamate side chains in ciliary motility has been established through these studies, the functional significance of short glutamate side chains remains elusive.

In this study, we characterized *Chlamydomonas* mutants lacking the CCP1, CCP2, and CCP5 orthologues. Although long polyglutamate side chains in the axoneme remained largely unaffected across all mutants, short glutamate side chains were significantly increased in the *ccp5-1* mutant axoneme. Strikingly, the *ccp5-1* mutation increased the swimming motilities in the background of *tpg1*. These findings suggest that short glutamate side chains – most likely monoglutamylation – is sufficient to modulate ciliary motility.

## Materials and Methods

### Strains and Cultures

*Chlamydomonas reinhardtii* strains (Table 1) were maintained on tris-acetate-phosphate (TAP) agar plates (Gorman and Levine, 1965) for long term storage. For experiments, cells were cultured in liquid TAP medium at 25°C under a light/dark cycle of 12:12 h and aeration. Cell growth was monitored by measuring the optical density at 750 nm (*OD*750) at 24-hour intervals using a spectrophotometer.

**TABLE 1:** List of strains used in this study.

| Name | Origin | Mating type | Antibiotic resistance | Reference |
| --- | --- | --- | --- | --- |
| CC-124 |  | Minus |  | Pröschold et al (2005) |
| *tpg1* | CC5245 | Minus |  | Kubo et al (2010) |
| *ccp1-1* | CC-124 | Minus | Hygromycin | This study |
| *ccp1-1 tpg1* | CC-5245 | Minus | Hygromycin | This study |
| *ccp2-1* | CC-124 | Minus | Paromomycin | This study |
| *ccp2-1 tpg1* | CC-5245 | Minus | Paromomycin | This study |
| *ccp5-1* | CC-124 | Minus | Hygromycin | This study |
| *ccp5-1 tpg1* | CC-5245 | Minus | Hygromycin | This study |

\*Generated mutants will be deposited to the *Chlamydomonas* Resource Center (https://www.chlamycollection.org/) after publication.

### Mutant Generation

*Chlamydomonas* mutants were generated via CRISPR/Cas9-mediated gene editing, as previously described (Kubo et al., 2024). Briefly, target sequences were determined using CRISPRdirect (http://crispr.dbcls.jp/). Guide RNAs (gRNAs) were prepared by annealing specific crRNAs (Table 2) and tracrRNA (FASMAC, Kanagawa, Japan), followed by incubation with Cas9 protein (FASMAC) to form Cas9/gRNA ribonucleoprotein (RNP) complexes. For transformation, cells were pre-treated with autolysin to remove the cell wall (Picariello et al., 2020) and then electroporated with the Cas9/gRNA RNPs along with donor DNA using an electroporator (BTX ECM630; BXD). The donor DNA contained either hygromycin or paromomycin resistance cassettes flanked by homology arms. Following electroporation, cells were recovered in TAP sucrose medium and then plated on TAP-agar containing 10 μg/ml hygromycin or paromomycin. Resulting colonies were screened and confirmed by PCR using specific primers.

**TABLE 2:**
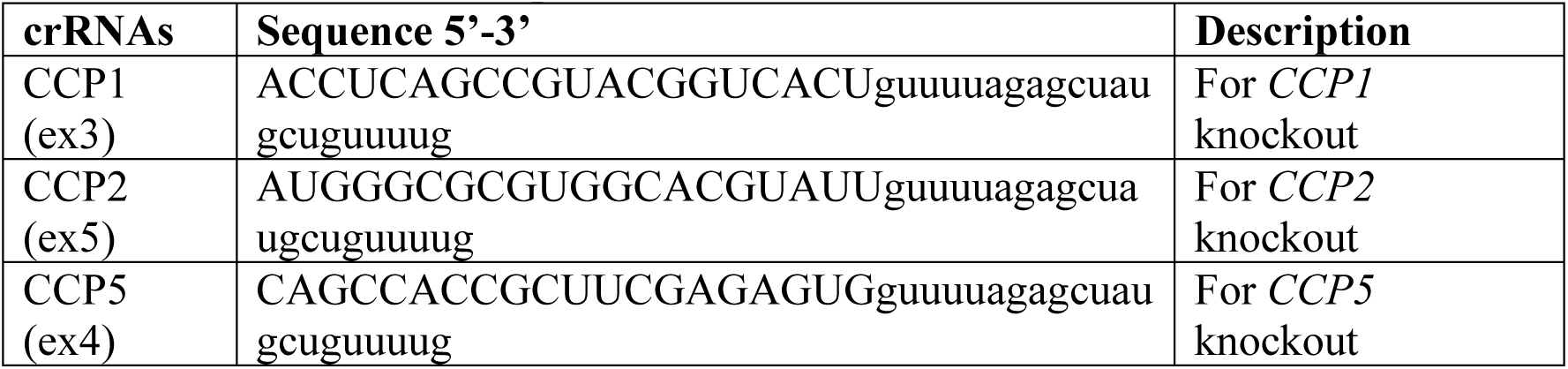
List of crRNA sequences.

### Reverse transcription quantitative PCR (RT-qPCR)

Total RNA was extracted from wild type, *ccp1-1*, *ccp2-1*, and *ccp5-1* mutants using Trizol LS (Molecular Research Center) according to the manufacturer’s instructions. 20 μl of cDNA was synthesized from 1 μg of total RNA using ReverTraAce (TOYOBO). Quantitative PCR was performed using SYBR Green qPCR Master Mix (ThermoFisher) with specific primer sets (Table 3) on a StepOnePlus system (ThermoFisher). The transcript levels of CCP1, CCP2, and CCP5 were normalized to the expression of G protein subunit-like protein (GBLP; encoded by Cre06.g278222), and relative expression was calculated using the 2^-ΔΔ*Ct*^ method.

**TABLE 3:**
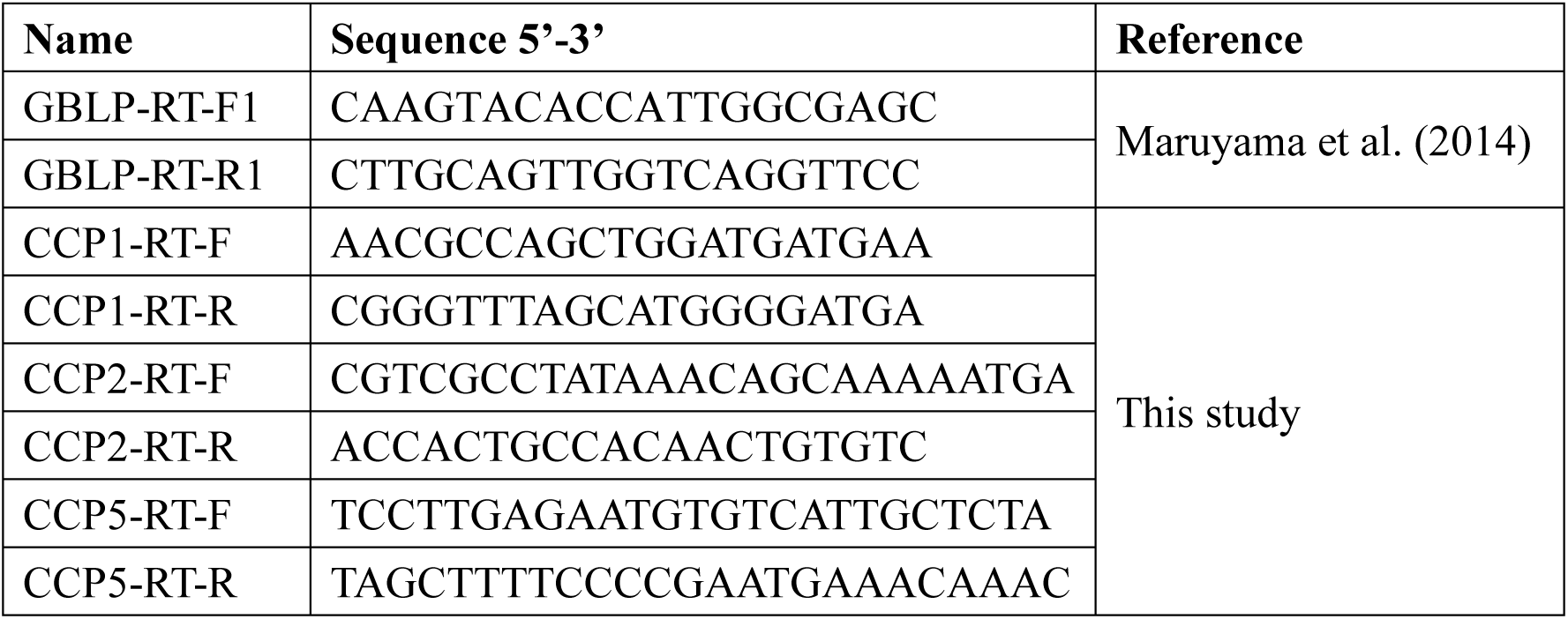
List of primers for RT-qPCR.

### Indirect immunofluorescence microscopy

Immunostaining of NFAps was carried out as previously described (Kubo et al., 2024; Sanders and Salisbury, 1995). Fully grown cells were suspended into 6 ml autolysin and gently agitated for 60 min to remove cell walls. The cells were washed with NB buffer (6.7mM Tris-HCl [pH7.2], 3.7mM EGTA, 10mM MgCl2, and 0.25mM KCl) and placed on an eight well-slide glass (8-mm well; Matsunami) treated with polyethyleneimine.

The cells were demembranated by 1% Igepal CA-630 (Sigma) and subsequently fixed with 2% paraformaldehyde in NB buffer for 10 min. The cells were then treated with acetone at -20℃ and methanol at -20℃ for 5 min each. The cells were first treated with blocking buffer (1% bovine serum albumin and 3% Fish Skin gelatin in phosphate-buffered saline [PBS]) and incubated with primary antibodies (Table 4) followed by secondary antibodies (goat anti-rabbit IgG Alexa 488, 1:200, Invitrogen; goat anti-mouse IgG Alexa Fluor 594, 1:200, Invitrogen). The cells were treated with antifade mountant (SlowFade Diamond, Thermo Fisher Scientific) and encapsulated with a glass coverslip. The sample was examined with a microscope (BX53, Olympus) and images were acquired with a CCD camera (ORCA-Flash4.0sCMOS, Hamamatsu).

**TABLE 4:** Antibodies used in this study.

| Antibody (clone) | Host | Dilution | Reference / Origin |
| --- | --- | --- | --- |
| Anti-acetylated tubulin (6-11B-1) | Mouse | 1:5000 (WB)<br>1:200 (IFM) | Sigma-Aldrich |
| Anti-tyrosinated tubulin (TUB-1A2) | Mouse | 1:3000 (WB)<br>1:200 (IFM) | Sigma-Aldrich |
| Anti-detyrosinated tubulin (AB3201) | Rabbit | 1:2000 (WB)<br>1:200 (IFM) | Merck |
| Anti-polyglutamylated tubulin (polyE#1) | Rabbit | 1:2000 (WB),<br>1:200 (IFM) | Kubo et al. (2017) |
| Anti-glutamylated tubulin (GT335) | Mouse | 1:2000 (WB),<br>1:200 (IFM) | Adipogen Life Sciences |
| Anti-α-tubulin (B512) | Mouse | 1:5000 (WB)<br>1:500 (IFM) | Sigma-Aldrich |

### Isolation of axoneme

Cilia were isolated following the method described in Kubo et al. (2024) according to a modified version of Witman et al. (1972). Briefly, fully grown cells were collected with centrifugation (3,000 rpm, 5 min) and treated with 1 mM dibucaine-HCl (Wako) to amputate their cilia. Cilia were washed and collected by centrifugation (15,000 rpm, 20 min, 4℃). Isolated cilia were suspended into HMDEK buffer (30 mM HEPES, 5 mM MgSO4, 1 mM DTT, 0.1 mM EGTA, and 25 mM CH3COOK) and demembranated by 0.1% Igepal CA-630 (Sigma) to prepare axonemal samples. For the immunofluorescence microscopy of axonemal microtubules, isolated axonemes were placed on eight-well slide glass and treated with 0.1 mM ATP (Sigma) and 0.5 μg/ml nagarse (Proteinase, bacterial Type XXIV; Sigma) to induce splaying. Immunostaining of the splayed axonemes was processed as described above in the “Indirect immunofluorescence microscopy” section.

### Assessment of motility

Swimming velocity was measured by tracking images of the moving cells as described in Kubo et al. (2024). Briefly, the cells under the dark-field microscope equipped with 40× objective were recorded using a digital camera with a frame rate of 30 fps, and the obtained movies were processed with ImageJ.

The ciliary beat frequency was measured according to Kamiya. (2000) with modification. Briefly, light intensity fluctuations in the microscopic images of swimming cells were captured using a custom-built detection unit equipped with a collecting lens and a high-gain current-to-voltage (I/V) converter, which was mounted directly onto the microscope tube. The resulting voltage signals were digitized and analyzed via fast Fourier transform (FFT) using the WaveSpectra software (developed by efu; https://efu.sub.jp/soft/ws/ws.html) to generate power spectra. For each strain, the dominant frequency peak was identified from the power spectrum, and this measurement was repeated 20 times to ensure data reproducibility. The mean beat frequency and standard deviation (SD) were then calculated from these 20 independent peak values.

## RESULTS

### Generation of *Chlamydomonas* mutants deficient in cytosolic carboxypeptidase genes (*CCP1*, *CCP2*, and *CCP5*) using CRISPR/Cas9-mediated gene editing

In mammals, the cytosolic carboxypeptidase (CCP) family comprises six members (CCP1-6) belonging to the M14D subfamily of metallocarboxypeptidases (Wu et al., 2014). A genome-wide search for deglutamylase orthologues in *Chlamydomonas reinhardtii* identified four primary candidates. Among these, CCP1 (also known as FBB17; Cre13.g572850), CCP2 (Cre02.g102850), and CCP5 (Cre12.g528100) displayed amino acid identities of 41.3% (E-value 1e-104), 37.3% (E-value 4e-44), and 43.0% (E-value 5e-45) to their respective murine counterparts. As illustrated in Figure 1A, CCP1, CCP2, and CCP5 share a common architecture consisting of a catalytic peptidase M14-like domain and a CCP-specific N-terminal domain. This N-terminal domain is hypothesized to play regulatory roles or to assist the structural assembly of these metallocarboxypeptidases. In contrast to these three proteins, Cre13.g565600 exhibited a significantly lower degree of conservation, with only 23.4% identity to mouse CCP5 (E-value 1e-13). Based on these homology scores, we focused our subsequent analyses on the first three candidates and excluded Cre13.g565600 from this study.

**Figure 1.**
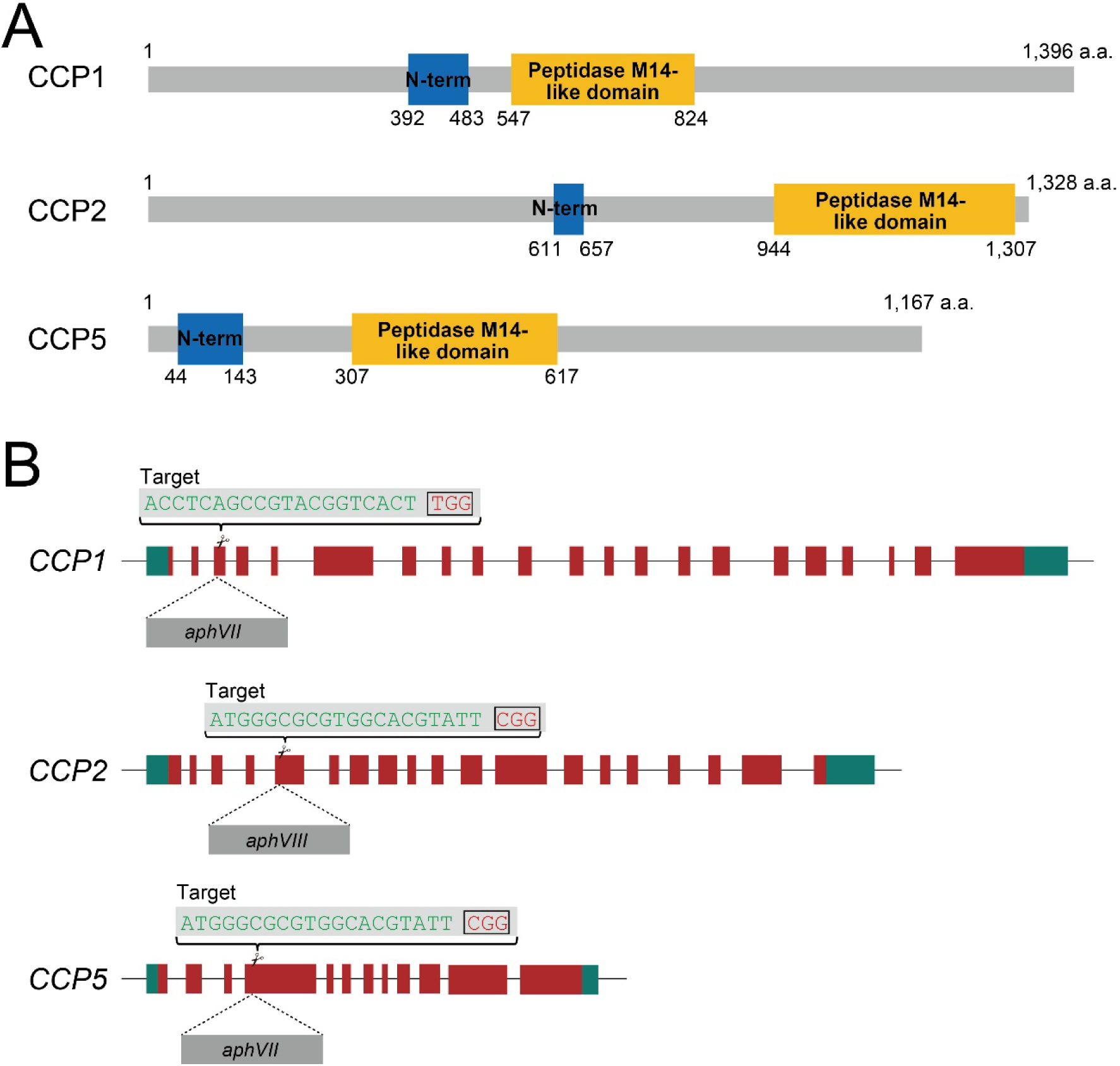
Generation of *ccp1-1*, *ccp2-1*, and *ccp5-1* mutants via CRISPR/Cas9-mediated gene editing. (A) Schematic protein structures of *Chlamydomonas* CCP1, CCP2, and CCP5. (B) Genomic organization of the *CCP1* (Cre13.g572850), *CCP2* (Cre02.g102850), and *CCP5* (Cre12.g528100) genes. Dark red boxes represent exons, and greenish blue boxes indicate untranslated regions (UTRs). To generate *ccp1-1*, *ccp2-1*, and *ccp5-1* mutants, a drug resistance cassette (dark gray; conferring resistance to either hygromycin or paromomycin) was inserted into the indicated target exons.

We generated mutants deficient in the genes encoding CCP1, CCP2, and CCP5 by CRISPR/Cas9-mediated gene editing. In the wild-type strain, hygromycin (*aphVII*) or paromomycin (*aphVIII*) resistant cassettes were introduced into the coding sequences of the *CCP1*, *CCP2*, and *CCP5* genes (Figure 1B). Transformants were then screened on medium containing the respective antibiotics. Genotyping using specific primer sets confirmed the site-specific integration of resistance cassettes, identifying several independent transformants for each gene. Specific clones deficient in the *CCP1*, *CCP2*, and *CCP5* genes were designated as *ccp1-1*, *ccp2-1*, and *ccp5-1*, respectively. Gene expression levels of these clones were analyzed by RT-qPCR. As expected, all clones exhibited significantly lower mRNA levels of the respective target genes compared to the wild type (Table 5). The marked reduction of mRNA suggests that aberrant transcripts may be degraded via nonsense mediated mRNA decay. Regardless, any potentially truncated products are likely non-functional as evidenced by the subsequent defects in tubulin modification described below.

**TABLE 5:**
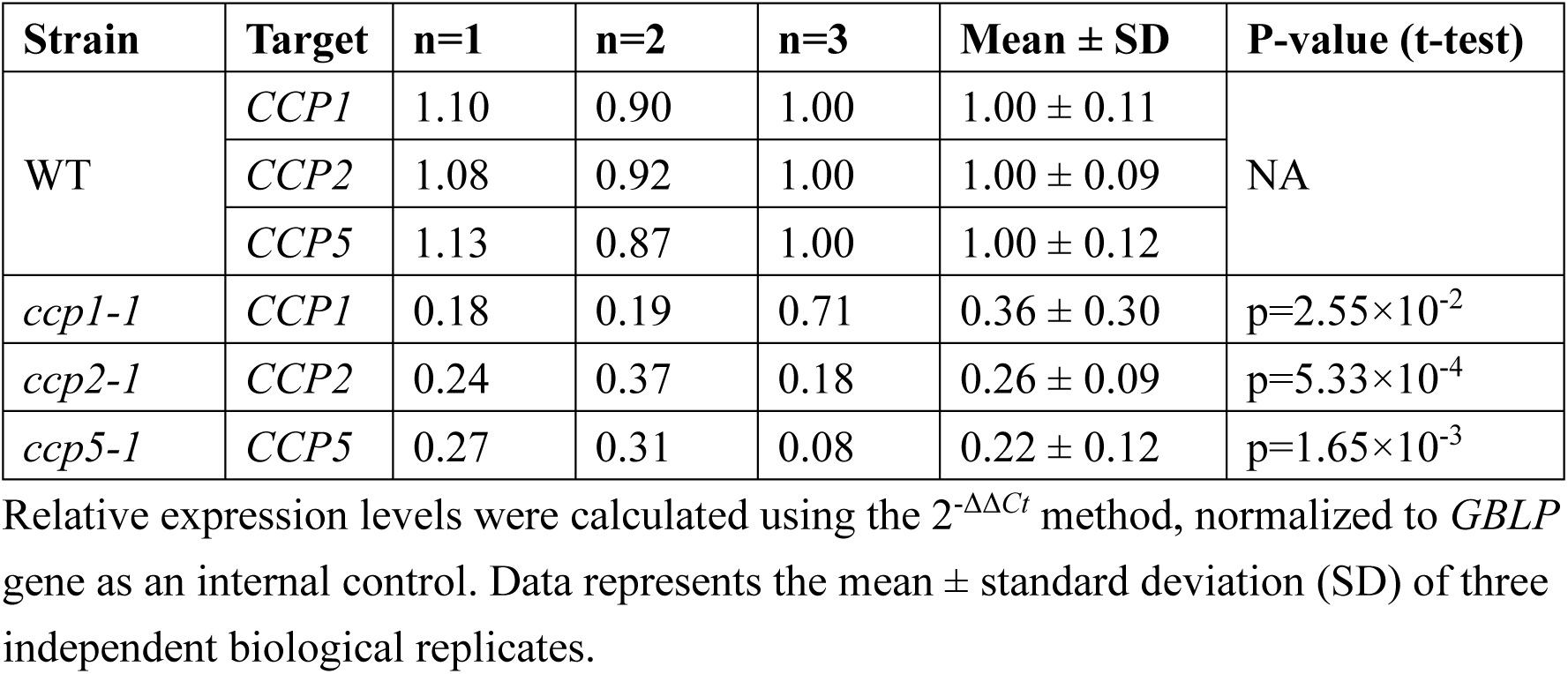
RT-qPCR analysis of *CCP1*, *CCP2*, and *CCP5* gene expression.

| Strain | Target | n=1 | n=2 | n=3 | Mean ± SD | P-value (t-test) |
| --- | --- | --- | --- | --- | --- | --- |
| WT | *CCP1* | 1.10 | 0.90 | 1.00 | 1.00 ± 0.11 | NA |
| | *CCP2* | 1.08 | 0.92 | 1.00 | 1.00 ± 0.09 | |
| | *CCP5* | 1.13 | 0.87 | 1.00 | 1.00 ± 0.12 | |
| *ccp1-1* | *CCP1* | 0.18 | 0.19 | 0.71 | 0.36 ± 0.30 | p=2.55×10<sup>-2</sup> |
| *ccp2-1* | *CCP2* | 0.24 | 0.37 | 0.18 | 0.26 ± 0.09 | p=5.33×10<sup>-4</sup> |
| *ccp5-1* | *CCP5* | 0.27 | 0.31 | 0.08 | 0.22 ± 0.12 | p=1.65×10<sup>-3</sup> |

Relative expression levels were calculated using the $2^{-\Delta\Delta C_t}$ method, normalized to *GBLP* gene as an internal control. Data represents the mean ± standard deviation (SD) of three independent biological replicates.

### Loss of any CCP has minimal impact on axonemal tubulin polyglutamylation

The levels of polyglutamylated tubulin in these mutants were examined by immunofluorescence microscopy. Nuclear-flagellar apparatuses (NFAps) were isolated and double-stained for polyglutamylated tubulin (Kubo et al., 2017; polyE) and acetylated-α-tubulin. Unexpectedly, axonemal polyglutamylation levels in all mutants were comparable to those of the wild type (Figure 2A).

**Figure 2.**
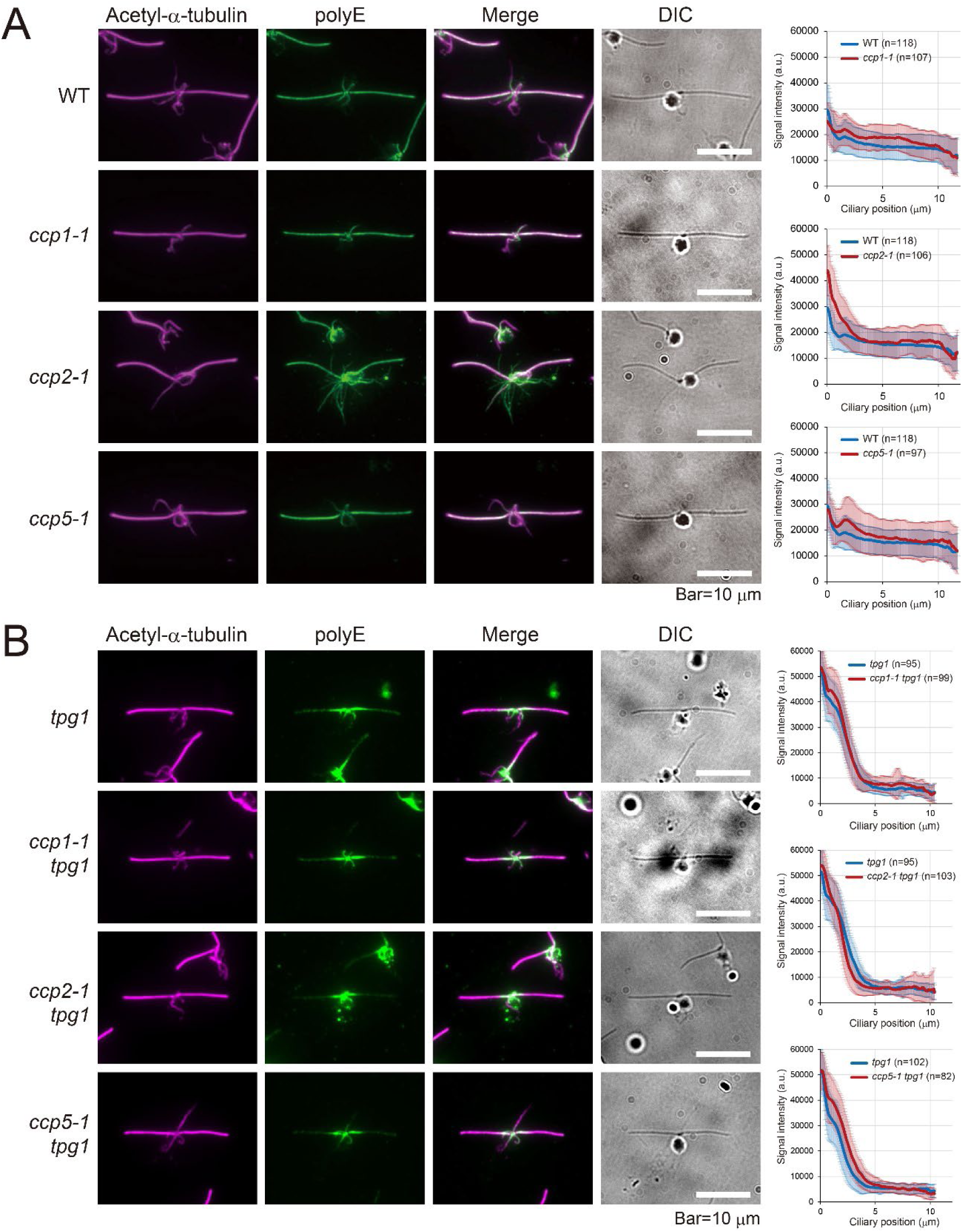
Axonemal polyglutamylation levels are normal in *ccp* mutants. Localization of polyglutamylated tubulin in *ccp* mutants. (A, B) Immunofluorescence images of *ccp1-1*, *ccp2-1*, and *ccp5-1* NFAps in either a wild-type background (A) or a *tpg1-*deficient background (B). NFAps were double-stained with antibodies against acetylated α-tubulin (as a structural marker) and polyglutamylated tubulin (polyE).

The polyE signals were further evaluated in a *tpg1* background. The *tpg1* mutant lacking the elongase TTLL9 is known to have reduced distal staining in the axoneme when stained with polyE antibody (Kubo et al., 2010; Figure 2B). While *ccp5-1 tpg1* showed a slight increase in the polyE staining, axonemal signals in all double mutants remained consistently low, with no drastic recovery from the *tpg1* staining pattern (Figure 2B).

### CCP5 deficiency increases short glutamate chains in the axonemal tubulin

Next, we assessed glutamylation using the GT335 antibody, which recognizes the branch-point glutamate independently of the chain length. In general, GT335 labels the entire length of the axonemes, with the signal intensity decreasing towards the tip (Figure 3). While *ccp1-1* and *ccp2-1* axonemes showed minimal or no increase, *ccp5-1* axoneme exhibited a significant elevation in GT335 signal (Figure 3A). Combined with its subtle increase in polyE (Figure 2), this result strongly suggests that CCP5 primarily deglutamylates the branch-point glutamate instead of shortening polyglutamate chains.

**Figure 3.**
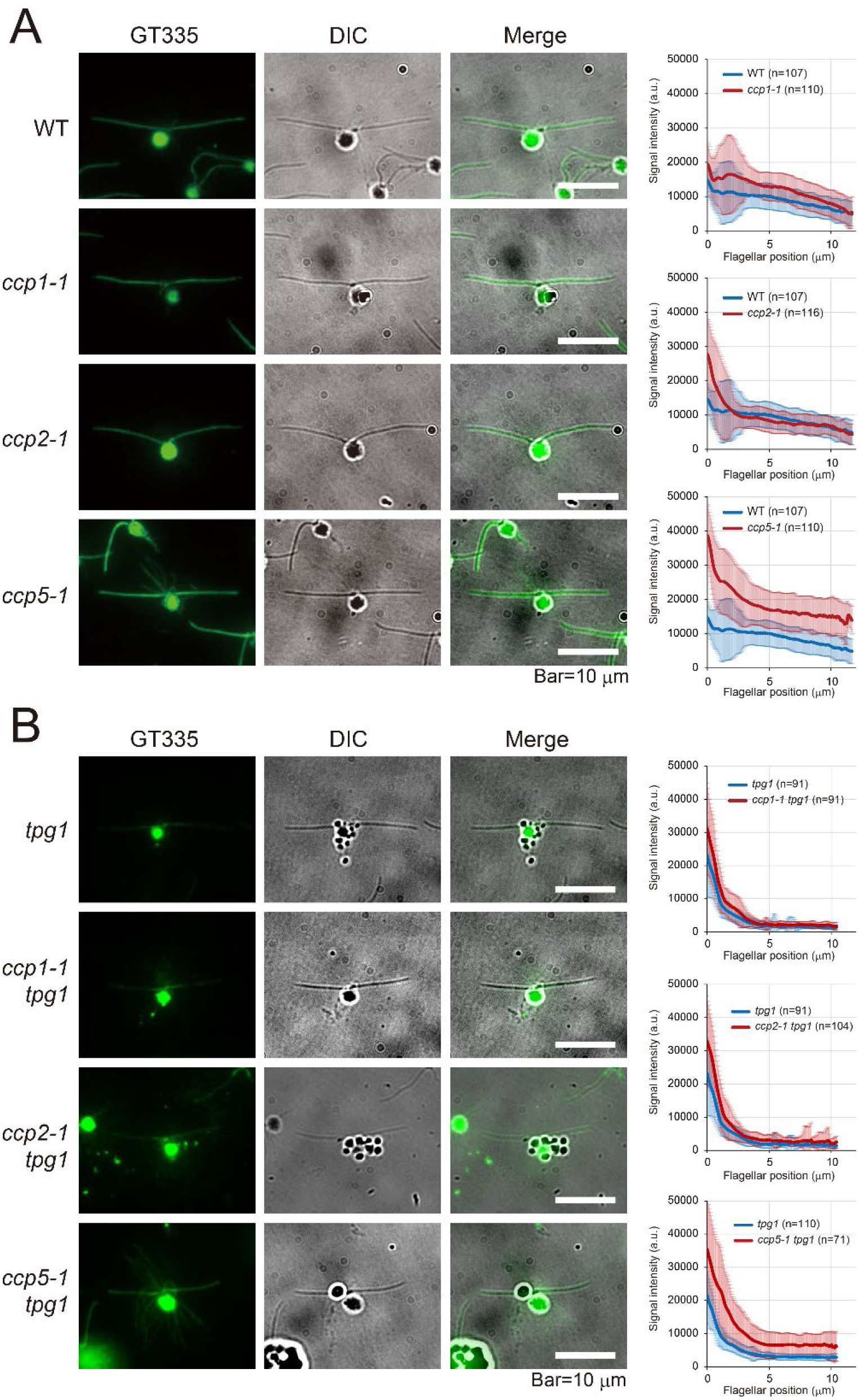
Accumulation of monoglutamylated tubulin in the *ccp5-1* mutant axoneme. Localization of branch point glutamylation in *ccp* mutants. (A, B) Immunofluorescence and DIC images of *ccp1-1*, *ccp2-1*, and *ccp5-1* NFAps in either a wild-type background (A) or a *tpg1-*deficient background (B). NFAps were stained with the GT335 antibody, which specifically recognizes the glutamylation branch point (representing both mono-and polyglutamylation). Note the marked increase in GT335 signal in *ccp5-1* mutant axonemes in both backgrounds compared to the other strains.

The GT335 signals of *ccp* mutants were further examined in the *tpg1* background. Compared to the wild type, the *tpg1* single mutant exhibited reduced axonemal staining, indicating that glutamylation levels are generally low in this mutant. In *ccp1-1 tpg1* and *ccp2-1 tpg1*, the axonemal staining remained comparable to those of the *tpg1* single mutant (Figure 3B). In contrast, the *ccp5-1 tpg1* double mutant exhibited markedly increased signal in the axoneme (Figure 3B). Therefore, our findings suggest that CCP5 functions as a deglutamylase targeting short glutamate side chains or monoglutamylated tubulin in the axoneme.

### The lack of either CCP1 or CCP2 results in the accumulation of detyrosinated tubulin in the axoneme

While CCPs are primarily recognized for shortening glutamate side chains, they also act as tubulin carboxypeptidase that processes the gene-encoded C-terminal glutamates. Specifically, CCP1 and CCP2 cleave the C-terminal glutamate residue of detyrosinated tubulin to generate Δ2-tubulin (Rogowski et al., 2010; Berezniuk et al., 2012; 2013; Tort et al., 2014; Song and Brady, 2015), whereas CCP5 is suggested to catalyze further processing into Δ3-tubulin (Berezniuk et al., 2013). To investigate the carboxypeptidase activity of these enzymes, we examined detyrosinated tubulin levels in *Chlamydomonas* CCP mutants. In controls, detyrosination was detected throughout the axoneme but absent in the cytoplasm (Figure 4A, 4B). All three mutants in the wild-type background showed abnormally high detyrosination levels, though the increase in *ccp5-1* was modest compared to *ccp1-1* and *ccp2-1* (Figure 4A). These results suggest that CCP1 and CCP2 are the enzymes acting for detyrosinated tubulin processing, likely leading to Δ2-tubulin generation, while the contribution of CCP5 to Δ2-tubulin processing is relatively limited.

**Figure 4.**
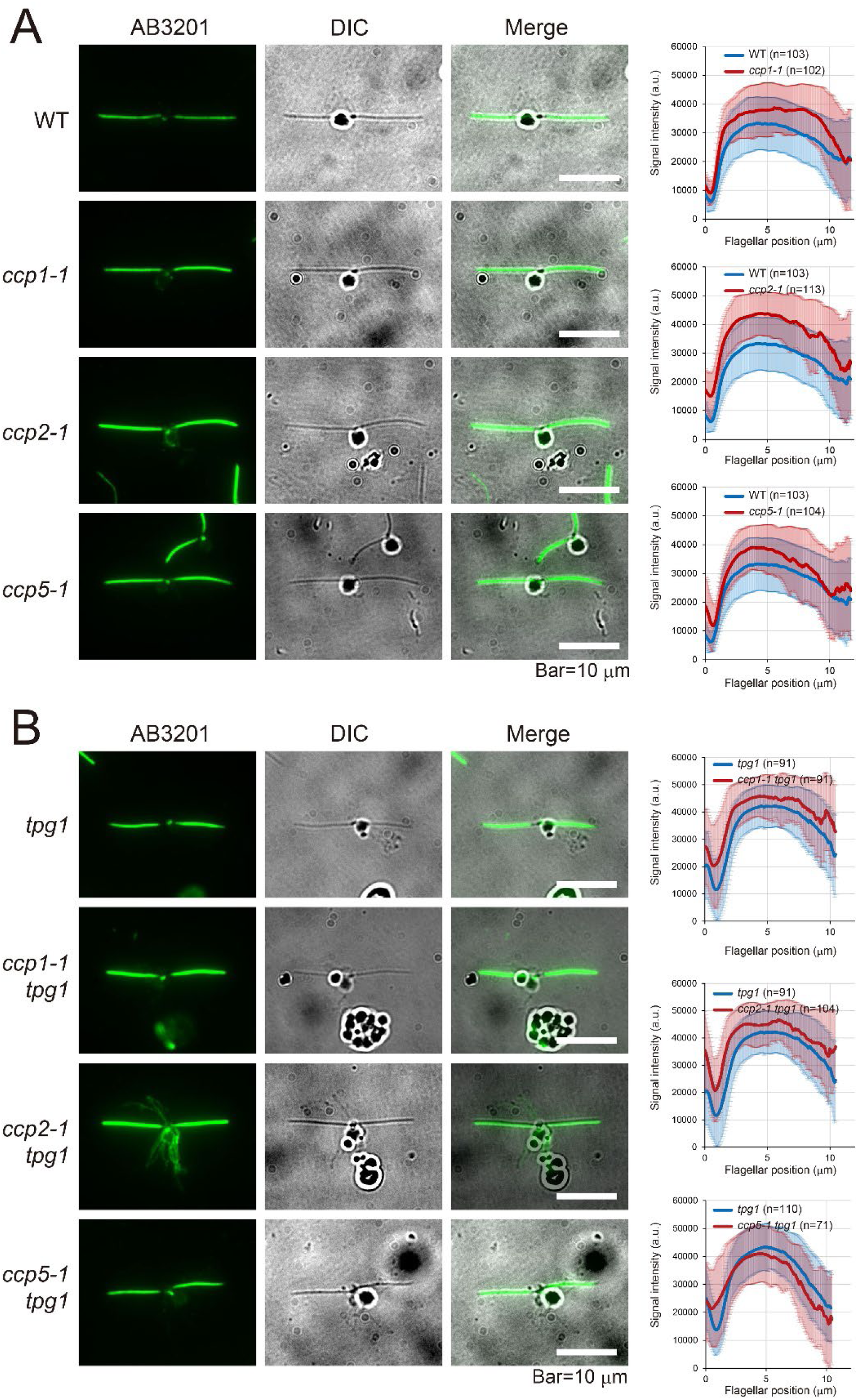
Accumulation of detyrosinated tubulin in *ccp* mutant axonemes. Localization of detyrosinated tubulin in *ccp* mutants. (A, B) Immunofluorescence and DIC images of *ccp1-1*, *ccp2-1*, and *ccp5-1* NFAps in either a wild-type background (A) or a *tpg1-*deficient background (B). NFAps were stained with an antibody against detyrosinated tubulin (AB3201).

This pattern was also observed in the *tpg1* mutant background, although the differences were narrower than in the wild type. Notably, *ccp5-1 tpg1* showed almost no difference from the *tpg1* single mutant (Figure 4B). These results suggest that carboxypeptidase activities of CCPs may be influenced by the level of polyglutamylation. In any case, our data support a model where CCP1 and CCP2 act as the major enzymes generating Δ2-tubulin, while CCP5 facilitates further processing into Δ3-tubulin.

### The lack of CCP changes the modification levels of cortical microtubules

During our immunofluorescence analysis, we observed altered staining patterns in the cytoplasmic microtubules, particularly in the *ccp2-1* and *ccp5-1* mutants. To better visualize these changes, images were processed with an ImageJ edge-detection filter, which enhances the contrast by highlighting sharp gradients in pixel intensity.

In *Chlamydomonas*, acetylated α-tubulin is detected exclusively in the rootlet microtubules, which consist of four stable microtubule bundles in the cell body (LeDizet and Piperno, 1986). We found that the localization of polyglutamylated tubulin generally coincided with these acetylated signals, as confirmed by the double-staining (Figure 2A). While the *ccp1-1* and *ccp5-1* mutants showed normal polyE staining restricted to the rootlet microtubules, the *ccp2-1* mutant exhibited a marked accumulation of polyglutamylated tubulin in the cytoplasmic cortical microtubules (Figure 2A, 5A, 5B). This change suggests that CCP2 primarily functions to regulate polyglutamylation state within the cell body. Notably, this ectopic accumulation was completely abolished in the *ccp2-1 tpg1* double mutant (Figure 2B). This finding indicates a functional interplay between TTLL9 and CCP2 in the cytoplasm, where TTLL9-mediated polyglutamylation is counteracted by CCP2-dependent deglutamylation.

We next examined the staining patterns of the GT335 antibody. Remarkably, GT335 labeled four stable microtubule bundles in wild-type cells, which we identified as the rootlet microtubules based on their characteristic morphology (Figure 5A). To our knowledge, this is the first report of GT335-positive signals localized to the rootlet microtubules in *Chlamydomonas*. While this pattern was typical for most strains, the *ccp5-1* mutant uniquely exhibited an extensive accumulation of GT335 signals in the cytoplasmic cortical microtubules (Figure 5B). These results suggest that CCP5 specifically regulates branch-point glutamylation also in the cytoplasm.

**Figure 5.**
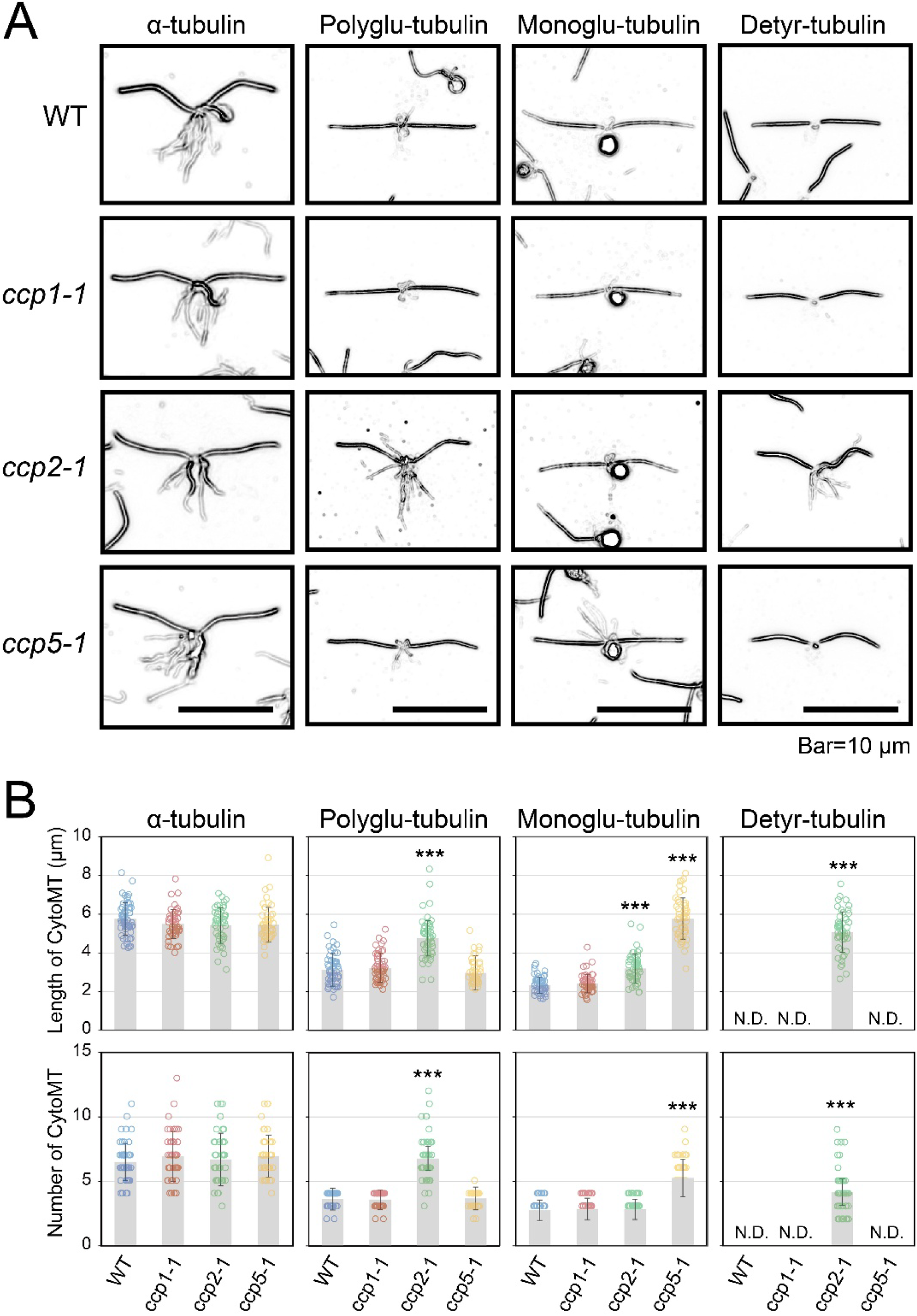
Accumulation of modified tubulins in the *ccp* mutant cytoplasm. (A) Immunofluorescence microscopy of NFAps using antibodies recognizing α-tubulin (B512), polyglutamylated tubulin (polyE#1), branch-point glutamate (GT335), and detyrosinated tubulin (AB3201). To better resolve fine cytoplasmic microtubule structures, an edge-detection filter was applied to the images using ImageJ. (B) Average length (upper panels) and number (lower panels) of stained cytoplasmic microtubules. The analysis was performed on approximately 50 cells. Values represent SD. Asterisks indicate significant differences compared with the controls (***p<0.001, Student’s t-test).

Finally, we examined the levels of detyrosinated tubulin in the cell body. In wild-type cells, the anti-detyrosinated tubulin antibody labeled only the axonemes and showed no detectable signal on any cytoplasmic microtubules (Figure 5A). Intriguingly, however, the *ccp2-1* mutant exhibited ectopic staining of the cytoplasmic cortical microtubules with this antibody. This result indicates that the loss of CCP2 leads to the accumulation of not only polyglutamylated but also detyrosinated tubulin in cytoplasmic microtubules, suggesting a broader role for CCP2 in maintaining the modification profile of the cortical cytoskeleton. Taken together, the immunofluorescence images obtained thus far indicate that (i) CCP1 primarily functions as a tubulin carboxypeptidase within cilia, (ii) CCP2 acts as a deglutamylase in the cytoplasm while serving as a tubulin carboxypeptidase both in cilia and cytoplasm, and (iii) CCP5 functions as a branch-point deglutamylase in both cilia and cytoplasm. These functional localizations and enzymatic activities of CCP enzymes are schematically summarized in Figure 6A and 6B.

**Figure 6.**
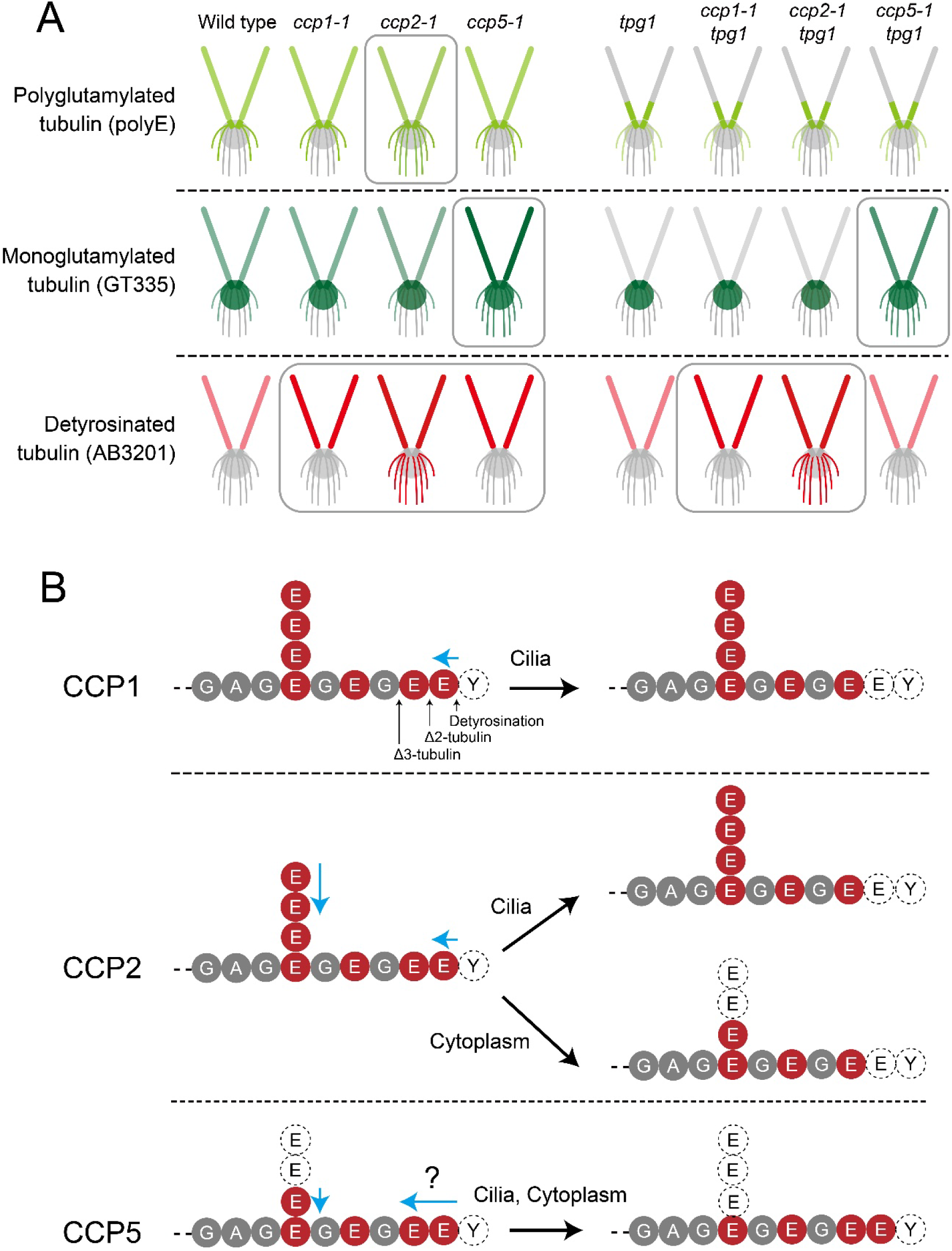
Summary of enzymatic activity of CCP1, CCP2, and CCP5. (A) Summary of tubulin modification defects in *ccp* mutants. polyE: Accumulation in *ccp2-1* cytoplasmic microtubules (absent in *ccp2-1 tpg1*). GT335: Accumulation in *ccp5-1* axonemes and cytoplasmic microtubules. AB3201: Accumulation in axonemes of *ccp1-1*, and *ccp2-1*; *ccp2-1* also shows abnormal cytoplasmic microtubule staining. (B) Summary of the suggested activities of CCP1, CCP2, and CCP5 in the cilia and cytoplasm. C-terminal sequences of α-tubulin are shown. Light blue arrows indicate either the deglutamylation or carboxypeptidase suggested by the *ccp* mutants.

**Figure 7.**
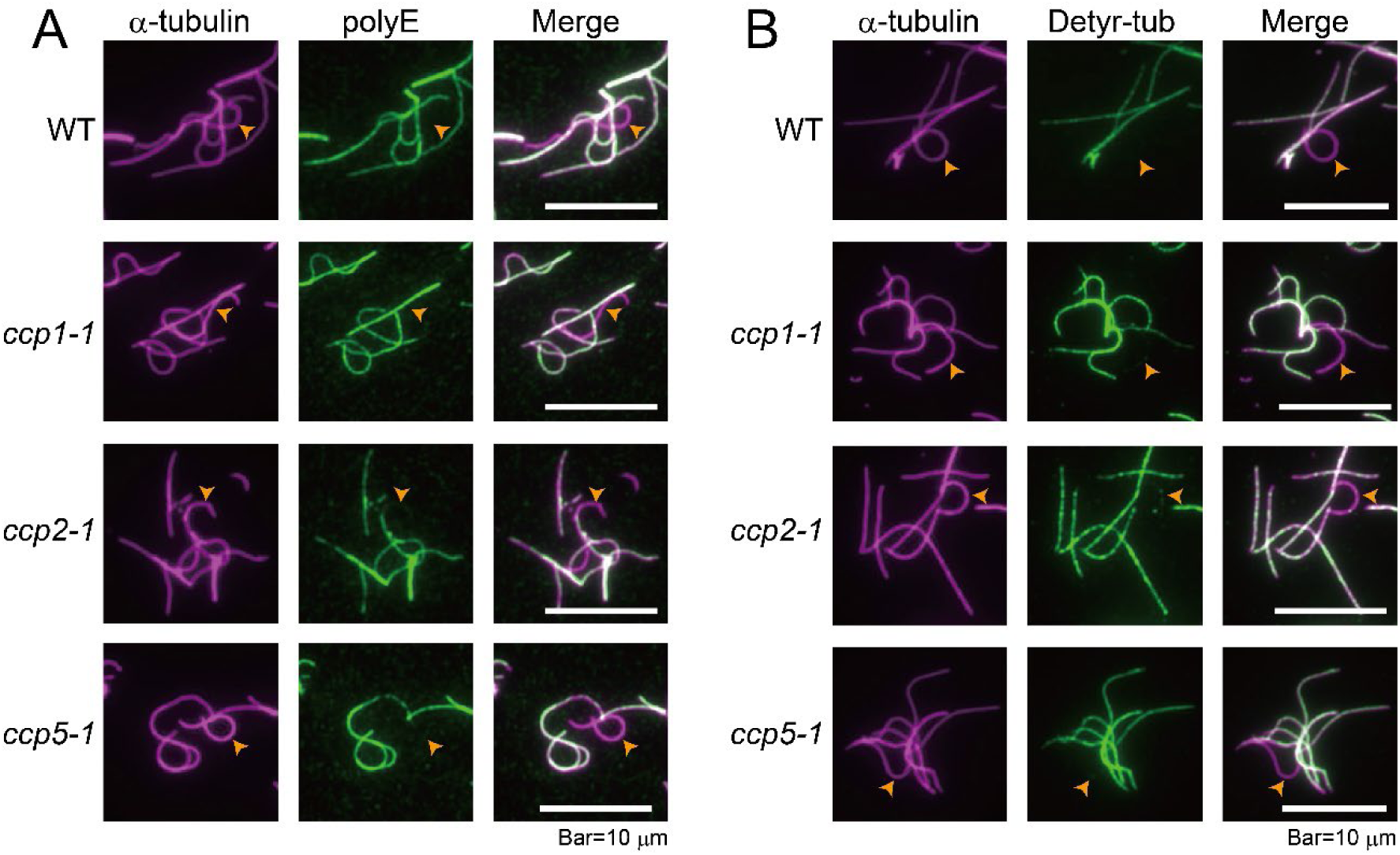
Absence of tubulin modifications in the central-pair microtubules of *ccp* mutants. Localization of polyglutamylated (A) and detyrosinated tubulin (B) in disintegrated axonemes. In all examined *ccp* mutants, both polyglutamylation and detyrosination signals were specifically absent from the central-pair microtubule, whereas they remained abundant in the outer-doublet microtubules. This observation suggests that tubulin-modifying enzymes, such as TTLL9, are specifically localized to the outer doublet microtubules and are absent from the central-pair complex.

### The central-pair microtubules remain unmodified in the absence of CCPs

The A- and B-tubules of the outer-doublet microtubules as well as the central-pair microtubules are composed of tubulins with distinct modification patterns (Multigner et al., 1996; Johnson, 1998; Lechtreck and Geimer, 2000). Notably, polyglutamylated tubulin is absent from the central-pair microtubules in *Chlamydomonas* (Lechtreck and Geimer, 2000; Kubo et al., 2010). This suggests that polyglutamylases, such as axonemal TTLL9, function exclusively on the outer-doublet microtubules, or that deglutamylases function to remove the polyglutamate side chains from the central-pair microtubules. To test this, we performed immunostaining of disintegrated axonemes from wild-type, *ccp1-1*, *ccp2-1*, and *ccp5-1* strains. In the disintegrated axonemes, we observed that a single microtubule bundle tended to remain unlabeled by both anti-polyglutamylated and anti-detyrosinated tubulin antibodies, whereas the other microtubules were clearly labeled (Figure 5A, 5B). The unstained bundles were identified as the central-pair microtubules based on their characteristic highly curved morphology (Kamiya et al., 1982; Kubo et al., 2010). These results suggest that TTLL9 acts exclusively on the outer-doublet microtubules, and that the CCP proteins studied here do not contribute to the modification patterns of the central-pair microtubules.

### CCP5 deficiency rescues the motility defect in the *tpg1* mutant

To investigate functional impact of altered tubulin modifications, the swimming behaviors of *ccp* mutants were examined. Initial observations revealed that all *ccp* single mutants exhibited slightly reduced swimming velocities compared to the wild type (Figure 8A), suggesting that precise regulation of tubulin modification levels is essential for normal ciliary motility. While *ccp1-1* and *ccp2-1* displayed normal beat frequencies (Figure 8B), *ccp5-1* unexpectedly exhibited increased beat frequency (Figure 8B). These distinct tendencies were largely maintained in the *tpg1* background. Notably, while *ccp1-1 tpg1* and *ccp2-1 tpg1* double mutants displayed more pronounced motility defects than the *tpg1* single mutant (Figure 8A), *ccp5-1 tpg1* exhibited a significant recovery in both swimming velocity and beat frequency (Figure 8A, 8B).

**Figure 8.**
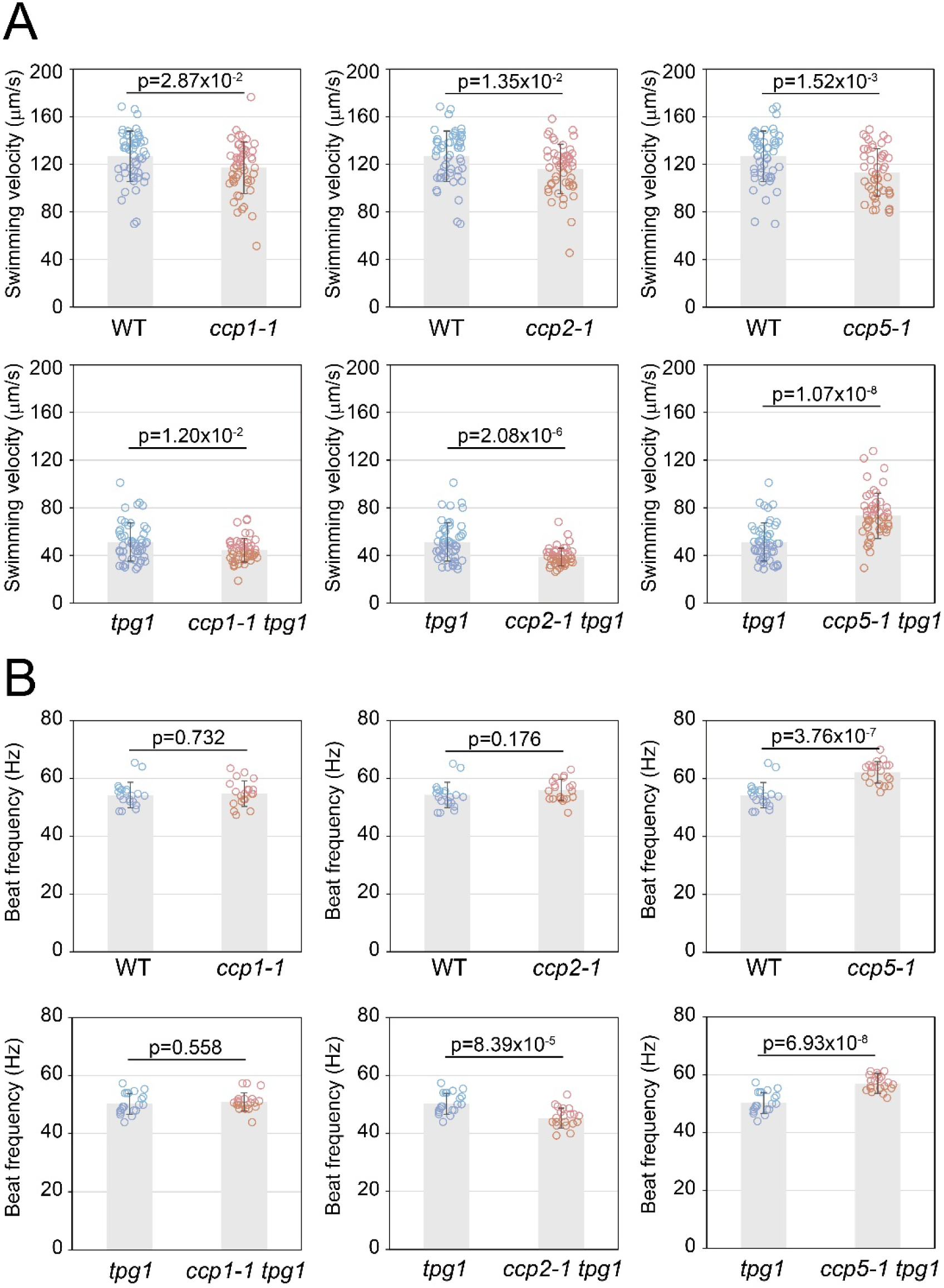
Synergistic enhancement of swimming motility in the *ccp5-1 tpg1* double mutant. (A) Swimming velocities and (B) beat frequencies of *ccp1-1*, *ccp2-1*, and *ccp5-1* strains in either a wild-type or a *tpg1*-deficient background. Note that while the *tpg1* mutation typically reduces motility, the additional loss of CCP5 exerts a synergistic effect, significantly increasing the swimming motility.

This specific rescue indicates that the loss of CCP5 specifically compensates for the motility impairment induced by the *tpg1* mutation.

## DISCUSSION

### Compartment-specific roles of CCP1 and CCP2 in deglutamylation and Δ2-tubulin generation

While mammals possess six CCP isoforms (CCP1-6) (Kalinina et al., 2007; Rogowski et al., 2010; Wu et al., 2014), we found that *Chlamydomonas* contains only three core candidates. This difference likely reflects evolutionary adaptation to distinct cellular requirements. The smaller CCP repertoire in *Chlamydomonas* may represent a simpler regulatory system compared to the specialized functions found in multicellular organisms. For instance, we identified orthologues of CCP1 and CCP2, which are known to remove glutamates from polyglutamate side chains and gene-encoded C-termini in other species (Rogowski et al., 2010; Berezniuk et al., 2012; Tort et al., 2014). Unexpectedly, however, our analysis using *ccp1-1* and *ccp2-1* mutants revealed that these enzymes have minimal effect on axonemal tubulin polyglutamylation. We however observed significant increase in detyrosinated tubulin levels within the axoneme of these two mutants. This result suggests that in the *Chlamydomonas* axoneme, CCP1 and CCP2 primarily function as enzymes responsible for the generation of Δ2-tubulin, rather than as deglutamylases that shorten polyglutamate side chains. It is possible that axonemal tubulin polyglutamylation, primarily mediated by TTLL9 (Kubo et al., 2010), is strictly maintained from excessive turnover to ensure stable ciliary motility.

In general, glutamylated tubulin in cytoplasm is restricted to the stable structure called the rootlet microtubules and is absent from the dynamic cortical microtubules (Figure 2, 6). Interestingly, however, polyglutamylated tubulin became detectable in the cortical microtubules of the *ccp2-1* mutant (Figure 2A). Furthermore, we found that this abnormal accumulation of polyglutamylated tubulin in *ccp2-1* is completely abolished in the *ccp2-1 tpg1* double mutant. These results indicate that CCP2 is responsible for removing polyglutamate side chains added by TTLL9 from cortical microtubules, thereby preventing their accumulation on these dynamic structures. Why cortical microtubules undergo deglutamylation, and how this process affects cellular homeostasis, remains an open question for future research.

### CCP5 is a universal deglutamylase that removes branch-point glutamates in both the axoneme and cytoplasm

Among the CCP family, only CCP5 is known to exhibit activity that removes the branch-point glutamates (Rogowski et al., 2010; Wu et al., 2017). Previous biochemical studies have shown that CCP5 is unique among the CCP family for its ability to remove the branch-point glutamates (Rogowski et al., 2010), and more importantly, it acts on both dimeric tubulin and polymerized microtubules (Berezniuk et al., 2013; Chen et al., 2024). In mammals and zebrafish, the lack of CCP5 orthologues impairs ciliary formation and function, leading to phenotypes such as sperm flagellar defects and photoreceptor disorganization (Wu et al., 2017; Pathak et al., 2014; Mercey et al., 2024). In contrast, the *ccpp-6* mutant of C. elegans shows accumulated glutamylated tubulin without severe structural defects (Kimura et al., 2010).

Consistent with the findings in *C. elegans*, our results demonstrate that the loss of CCP5 in *Chlamydomonas* leads to an increased GT335 signal intensity in both the axoneme and the cytoplasm without causing observable axonemal disorganization. This indicates that while CCP5 is essential for limiting tubulin glutamylation levels, it is not required for the fundamental assembly of the axoneme. Notably, despite the marked increase in the GT335 signal (Figure 3), the *ccp5-1* mutant exhibited near-normal levels of polyE signal (Figure 2). Since the GT335 antibody targets the branch-point glutamates, these data suggest that the loss of CCP5 primarily causes an accumulation of monoglutamylated tubulin rather than the elongation of existing side chains. Thus, the increased GT335 signal observed in both the ciliary axoneme and the cytoplasmic microtubules of *ccp5-1* indicates that CCP5 functions as a universal deglutamylase in *Chlamydomonas*.

### Role of monoglutamylation in modulating ciliary motility

A key finding of this study is that *ccp5-1* mutation partially rescues the motility defects of the *tpg1* mutant. Our results show that the *ccp5-1* mutation dramatically increases the GT335 signal even in the *tpg1* background (Figure 3B), indicating that CCP5 normally restricts the abundance of these short-chain species. While the basal level of glutamylation in *tpg1* is insufficient for normal dynein regulation, the increased frequency of these branch-point glutamates in *ccp5-1 tpg1* likely restores sufficient electrostatic interactions or structural docking sites required for the nexin-dynein regulatory complex (N-DRC) function (Kubo et al., 2012; 2017).

These findings redefine the functional role of tubulin modifications. Expanding on our previous work showing the necessity of polyglutamylation for N-DRC-mediated microtubule sliding (Kubo et al., 2012; 2017), the current results demonstrate that branch point monoglutamylation alone is sufficient to support N-DRC binding and partially restore ciliary motility. Thus, monoglutamylation is not merely a precursor for longer chains, but a distinct functional modification that independently modulates motor output. Mapping the precise axonemal localization of these branch-point glutamates will be a critical next step in understanding how they maintain ciliary integrity and mechanics.

## Acknowledgment

The authors acknowledge the use of Gemini (Google) for English language editing and proofreading during the drafting of this manuscript. This work was supported by Takeda Science Foundation (to T.K.), Institute for Fermentation, Osaka (to T.K.), The Kato Memorial Bioscience Foundation (to T.K.), Research Grant from Human Frontier Science Program [RGP006/2023 (to T.O.) (https://doi.org/10.52044/HFSP.RGP0062023.pc.gr.168592)], Japan Society for the Promotion of Science [23K05829 and 26K09375 (to T.K.), 21H02654 (to T.O.) and 24H02285 (to T.O.), and 26K01948 (to T.O.)].

